# Neuronal glutamate transporter EAAT3 regulates hippocampal GABAergic plasticity and reversal learning

**DOI:** 10.64898/2026.08.19.745605

**Authors:** Carlos Ancatén-González, Nicolás M. Ardiles, Sebastián F. Estay, Wladimir Plaza-Briceño, Alejandro Alcaíno, Pablo R. Moya, Andrés E. Chávez

## Abstract

Long-term depression (LTD) is a form of synaptic plasticity implicated in tasks involving the modification or elimination of previously learned information. While glial glutamate transporters can control the strength of synaptic plasticity, much less is known about the contribution of the neuronal glutamate transporter EAAT3 in controlling hippocampal LTD and learning processes. Here, we report that overexpression of EAAT3 in principal neurons, but not in GABAergic interneurons, impairs heterosynaptic GABAergic synaptic plasticity (iLTD) and homosynaptic excitatory LTD in the hippocampus. LTD impairments can be reversed by inhibiting EAAT3 or by a brief exogenous activation of mGluR during LTD induction, suggesting that, by limiting glutamate spillover between neighboring synapses, EAAT3 contributes to setting the strength of different forms of hippocampal LTD. Moreover, mice overexpressing EAAT3 in principal neurons, but not in GABAergic interneurons, display impaired reversal learning, a phenotype that can be rescued by blocking EAAT3 *in vivo*. Together, these findings reveal that, by controlling the strength of hippocampal LTD, EAAT3 contributes to cognitive flexibility required for processing new information.

**Significance statement:** Cognitive flexibility, particularly reversal learning, depends critically on the ability to weaken outdated synaptic connections, a cellular process mediated by long-term depression (LTD), that enables new memories to be stored in overlapping circuits. While astrocytic glutamate transporters are known to shape synaptic plasticity, the contribution of neuronal glutamate transporter EAAT3 has remained unclear. Here we identify EAAT3 as a key factor for hippocampal LTD and behavioral flexibility. Overexpression of EAAT3 in principal neurons, but not in GABAergic interneurons, impairs homosynaptic and heterosynaptic forms of LTD, and produces perseverative deficits in hippocampal-mediated reversal learning tasks that are rescued by EAAT3 blockade. These findings reveal that neuronal glutamate uptake, likely by limiting glutamate spillover between neighboring synapses, plays an important role in setting the threshold for multiple forms of hippocampal LTD and establishing a mechanistic link between EAAT3, synaptic depotentiation, and the capacity to update learned information *in vivo*.

## Introduction

Synaptic plasticity corresponds to the ability to change the neuronal connection strength - long-term potentiation (LTP) and long-term depression (LTD)- in an activity-dependent manner, and is considered part of the mechanisms underlying learning and memory processes(1). Although LTP has been widely studied as a physiological correlate of learning(1, 2), LTD has emerged as a critical and complementary form of synaptic plasticity in tasks involving the modification of previously learned information (3–13). Two major forms of homosynaptic LTD at excitatory synapses coexist in the hippocampus, including NMDA receptor–dependent and group I metabotropic glutamate receptor (mGluR)–dependent LTD (4, 14). Both forms of homosynaptic LTD have been shown to contribute to reversal learning, extinction, and cognitive flexibility, linking synaptic depotentiation to adaptive updating of behavior (3–6, 8, 10). In addition, hippocampal circuits also exhibit heterosynaptic LTD at inhibitory synapses (iLTD), where strong excitatory stimulation triggers mGluR activation and endocannabinoid release to reduce GABA release in a long-term manner (15, 16). Although a related form of iLTD has been linked to reversal learning and fear extinction (7), the mechanism that regulates iLTD and their contribution to cognitive flexibility remains poorly understood.

High-affinity membrane glutamate transporters (also named excitatory amino acid transporters, EAATs) are known to control the degree to which glutamate receptors could be activated during episodes of high neuronal activity and, thus, regulate the output of synaptic plasticity (17). EAAT type-3 (EAAT3), mostly found at postsynaptic sites (18, 19), limits glutamate escape between neighboring synapses and thereby constrains activation of NMDA receptors and mGluRs contributing to setting the strength of excitatory synaptic plasticity (20, 21). However, how EAAT3 contributes to the regulation of inhibitory synaptic plasticity and cognitive flexibility remains poorly understood. Interestingly, both findings in human brain tissue (22), and in animal models (23–25) suggest that increased EAAT3 expression could contribute to obsessive-compulsive disorder (OCD), a neuropsychiatric condition where impaired reversal learning has been reported (26–31). However, whether increased expression of EAAT3 impacts on the strength of hippocampal iLTD and hippocampal-dependent reversal learning *in vivo* remains unknown.

To address these issues, we assessed hippocampal synaptic function and plasticity in acute hippocampal slices and reversal learning in the Morris water maze in EAAT3 overexpressing mice. Our results indicate that overexpressing EAAT3 in pyramidal neurons does not affect basal excitatory and inhibitory synaptic transmission but impairs heterosynaptic GABAergic iLTD likely by limiting mGluRs activation. This deficit can be reversed by blocking EAAT3 and is absent when EAAT3 was overexpressed at inhibitory neurons. Finally, we found that overexpressing EAAT3 in principal neurons, but not in GABAergic interneurons, impaired reversal learning task, a phenotype that could be rescued by administering an EAAT3 blocker systemically. Altogether, these results suggest that EAAT3 might contribute to the regulation of the modification of previously learned information by controlling the strength of hippocampal LTD and establishing a mechanistic link between EAAT3, synaptic depotentiation, and the capacity to update learned information *in vivo*.

## Results

To study the impact of increased EAAT3 expression in controlling heterosynaptic GABAergic synaptic plasticity in mouse hippocampal slices, we first characterized basal excitatory synaptic transmission by monitoring spontaneous and miniature excitatory postsynaptic currents (sEPSCs and mEPSCs, respectively) from CA1 pyramidal neurons held at −60 mV (see methods) and found no differences in the amplitude or frequency of AMPAR-mediated sEPSCs and mEPSCs between EAAT3 overexpressing mice (EAAT3^glo^/CaMKII) and control littermates (EAAT3^glo^; **Figure 1A, Supplementary Figure 1A, Supplementary table 1**). Furthermore, no differences in the paired-pulse ratio (PPR) or short-term depression of evoked field excitatory postsynaptic potentials (fEPSPs) were observed between genotypes (**Supplementary Figure 1B, C; Supplementary table 1**). Moreover, the NMDAR/AMPAR ratio was similar between genotypes (**Figure 1B**), and no changes were observed in NMDAR current amplitudes when different levels of stimulation were delivered (**Supplementary Figure 1D, Supplementary table 1**). However, NMDAR-EPSCs showed slower decay kinetics in EAAT3^glo^/CaMKII mice compared with controls, suggesting an increase in expression of NR2B subunit (**Supplementary Figure 1E; Supplementary table 1**) that was reflected by a higher sensitivity to the NR2B antagonist Ro 25-6981 (5 μM) in EAAT3^glo^/CaMKII mice (**Supplementary Figure 1F; Supplementary table 1**). Altogether, these results indicate that, as previously demonstrated at cortical-striatal synapses(23), increased expression of EAAT3 at excitatory synapses does not affect basal or AMPAR-mediated synaptic transmission, but it increases the expression of the NR2B subunit of NMDARs at Schaffer collateral-to-CA1 synapses in the hippocampus.

**Figure 1.**
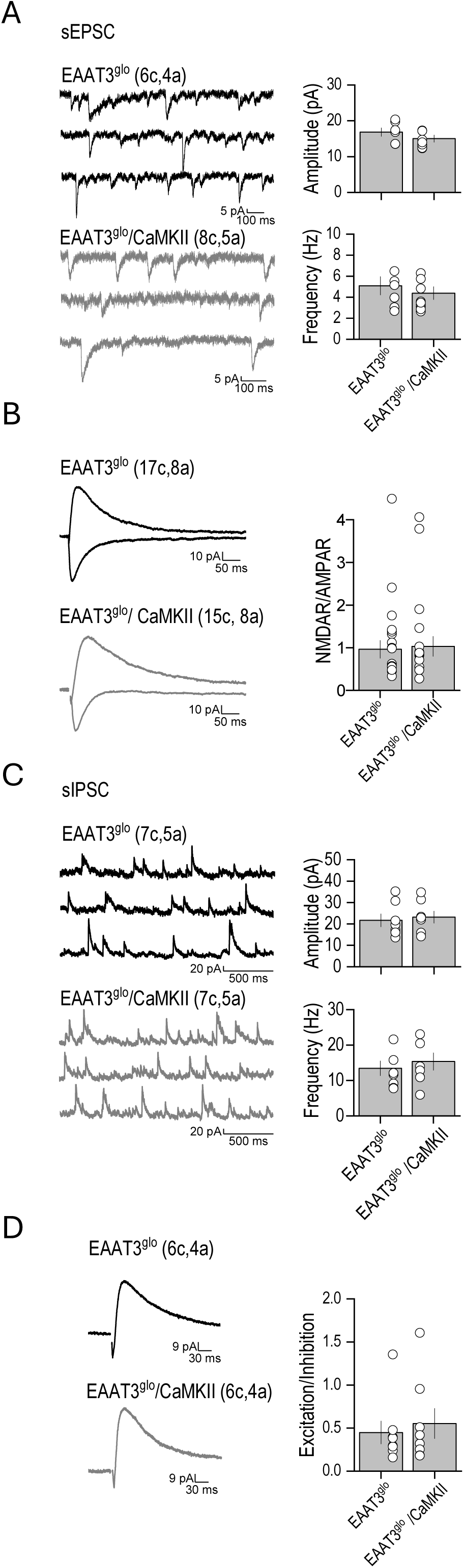
Overexpression of EAAT3 in principal neurons does not alter synaptic function in the CA1 area of the hippocampus. **A,** Spontaneous excitatory activity (sEPSC) is similar between EAAT3 overexpressing mice (EAAT3^glo^/CMKII) and control littermates (EAAT3^glo^). **B**, The AMPA/NMDA ratio is similar between genotypes. **C,** Spontaneous inhibitory activity (sIPSC) is also similar between genotypes. **D,** Excitatory (-60 mV) and inhibitory (-35 mV) responses measured from the same neuron display a similar ratio between genotypes, indicating that overexpression of EAAT3 does not alter basal synaptic function in the CA1 area of the hippocampus. In all panels, representative traces (left) and summarized plots (right) are shown. Data are presented as mean ± SEM, and the number of cells (c) and animals (a) are indicated in parenthesis.

Next, we characterized the basal inhibitory synaptic transmission by monitoring spontaneous and miniature inhibitory postsynaptic currents (sIPSCs and mIPSCs, respectively) from CA1 pyramidal neurons held at 0 mV (see methods) and found no differences in the amplitude or frequency of GABAR-mediated sIPSCs and mIPSCs (**Figure 1C; Supplementary Figure 2A**) between genotypes. Likewise, no differences were found in IPSCs paired-pulse ratio, short-term depression and IPSC amplitude input-output **(Supplementary Figure 2B-D**), indicating that EAAT3 overexpression at principal cells has no impact on basal hippocampal GABAergic synaptic function. Accordingly, when we simultaneously recorded evoked EPSC and IPSCs by holding pyramidal neurons at −35 mV and measured the excitatory/inhibitory balance at Schaffer collateral-to-CA1 synapses, no significant differences among genotypes were found (**Figure 1D**), indicating that EAAT3 overexpression does not alter the basal excitatory/inhibitory balance in the hippocampus.

As no changes in basal excitatory and inhibitory synaptic transmission were found (**Figure 1**), we next evaluated whether EAAT3 overexpression impacts glutamate spillover between neighboring synapses to regulate synaptic plasticity by monitoring a form of heterosynaptic inhibitory long-term depression (iLTD) that is independent of NMDAR but requires the activation of group I mGluR (15). Toward this end, we recorded CA1 pyramidal neurons while IPSCs were elicited by stimulating *stratum radiatum* fibers in the continuous presence of AMPA and NMDARs antagonists, NBQX (25 μM) and APV (25 μM) respectively. After a stable baseline, high-frequency stimulation (HFS; two trains of 100 stimuli at 100 Hz, separated by 20 s) induced robust GABAergic iLTD that was associated with a significant increase in PPR in control mice (**Figure 2A**), indicating that iLTD is due to a persistent reduction of presynaptic GABA release as previously reported (12). As iLTD induction requires activation of postsynaptic group I mGluR activation, we next evaluated whether EAAT3 overexpression can limit mGluR activation and thus impair iLTD. Consistent with this idea, HFS failed to induce iLTD and no changes in PPR were observed in EAAT3^glo^/CaMKII mice (**Figure 2A**). While EAAT3 expressed in GABAergic interneurons might strengthen inhibitory synapses in response to local increases in extracellular glutamate (32), we found that iLTD was normally induced in mice overexpressing EAAT3 in GABAergic cells (EAAT3^glo^/GAD) compared to their control littermates (EAAT3^glo^) (**Figure 2B**). Moreover, overexpressing EAAT3 in GABAergic cells does not alter GABAergic synaptic function as no differences between genotypes were found in sIPSCs, short-term depression, input-output curves and DHPG-induced iLTD, but a small, albeit, significative increase in both amplitude and frequency of mIPSCs, were detected in EAAT3^glo^/GAD mice (**Supplementary Figure 3, Supplementary Table 2**). These results strongly suggest that neuronal EAAT3 expressed at principal cells but not in GABAergic interneurons is important to regulate activity-dependent heterosynaptic plasticity at GABAergic synapses in the CA1 area of the hippocampus.

**Figure 2.**
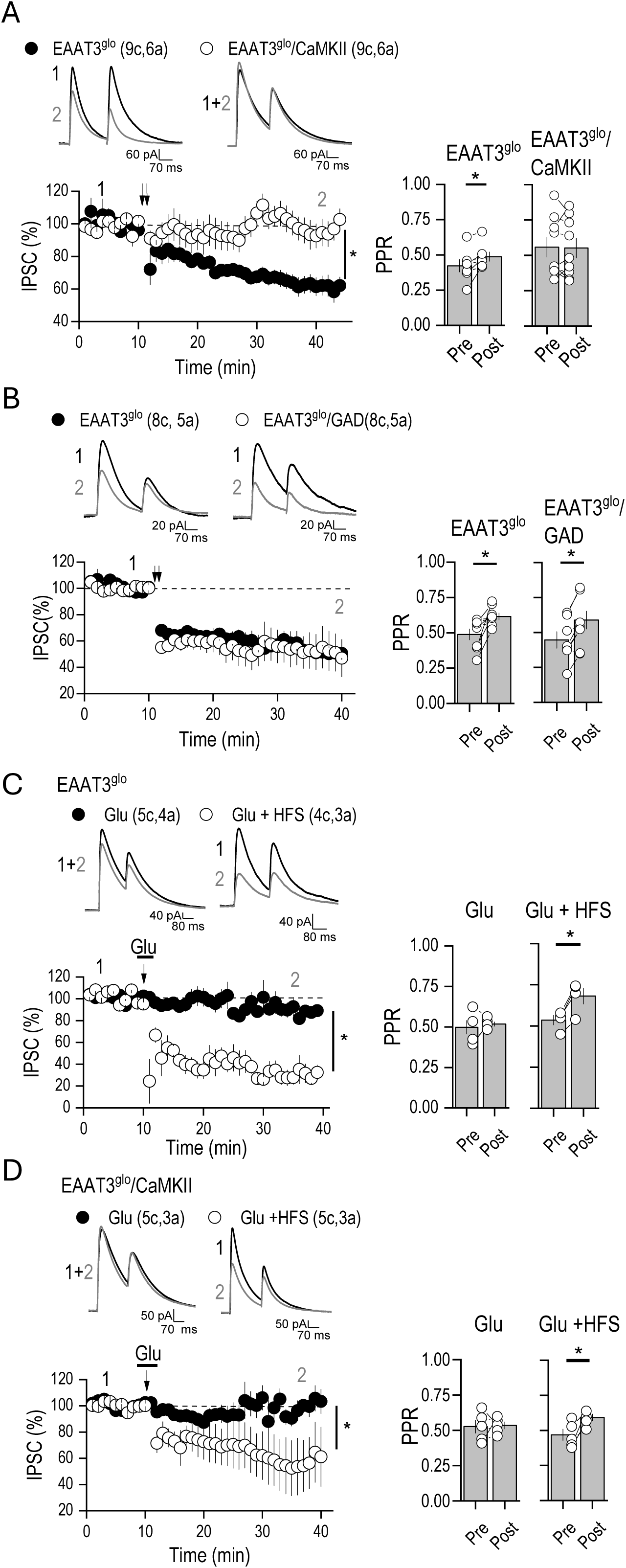
Increased EAAT3 expression in principal neurons, but not in GABAergic interneurons impair heterosynaptic GABAergic plasticity in the CA1 area of the hippocampus. **A,** Summary plots showing that two trains of high frequency stimulation (HFS; arrows) trigger a robust GABAergic iLTD that is associated with changes in paired pulse ratio (PPR) in control mice (EAAT3^glo^, black circles) but not in EAAT3 overexpressing mice (EAAT3^glo^/CMKII, while circles). **B**, HFS induces normal GABAergic iLTD in acute slices from animals overexpressing EAAT3 in GABAergic interneurons (EAAT3^glo^/GAD). **C,** Bath application of glutamate (Glu, 50mM) does not modify basal synaptic transmission at Schaffer collateral to CA1 synapses (black circles), but concomitant delivery of one train of 100 stimuli at 100 Hz, trigger a robust iLTD associated with changes in PPR in control mice. **D**, In slices from EAAT3^glo^/CaMKII mice, the application of glutamate together to one train at 100 Hz of stimulation induces iLTD, a synaptic depression that was accompanied by changes in PPR. In all panels, data are presented as mean ± SEM, and representative traces taken at times indicated by numbers are shown. The number of cells (c) and animals (a) are indicated in parenthesis. *P <0.05.

To test whether the impairment in the iLTD induction in EAAT3glo/CaMKII mice is due to a decrease in glutamate spillover by the increased expression of EAAT3, we first applied a brief exogenous application of glutamate (Glu, 50μM) to the bath in slices from control mice, that by itself was unable to modify basal synaptic transmission at Schaffer collateral to CA1 synapses (**Figure 2C**). However, when exogenous glutamate application was accompanied by one train of 100 stimuli at 100 Hz, which by itself also was unable to induce iLTD, a robust iLTD associated with changes in PPR was observed in control mice (**Figure 2C**). Importantly, in slices from EAAT3^glo^/CaMKII mice, the application of glutamate together to one train at 100 Hz of stimulation was able to induce iLTD that was accompanied by changes in PPR (**Figure 2D**). Under these experimental conditions a difference in the magnitude of iLTD was observed between control and EAAT3glo/CaMKII mice, supporting the importance of glutamate spillover in the size of inhibitory depression. Moreover, these results indicate that two HFS trains cannot produce sufficient glutamate spillover to activate mGluRs in EAAT3^glo^/CaMKII mice, but concomitant exogenous application of glutamate might saturate transporters allowing glutamate release during one train of high stimulation to escape to extrasynaptic sites to activate mGluRs and induce iLTD.

To confirm that the inability to induce iLTD in EAAT3^glo^/CaMKII mice is due to a reduction in glutamate spillover, next we tested the cellular mechanisms downstream to the glutamate spillover required to trigger iLTD. First, we tested whether mGluR function, required for the iLTD induction, is normal in EAAT3glo/CaMKII mice. To this end, we applied the mGluR agonist DHPG (50 μM) while monitoring GABAergic IPSCs. Bath application of DHPG was able to induce synaptic depression at same level in both genotypes (**Figures 3A**). DHPG-induced LTD was accompanied by changes in PPR indicating its presynaptic nature (**Figure 3A**). Moreover, we found that the level of group I mGluRs expression was similar between genotypes (**Figure 3B**), strongly suggesting that mGluRs are functional located to trigger synaptic depression in EAAT3^glo^/CaMKII mice. Likewise, application of the type 1 cannabinoid receptor (CB1R) agonist WIN55,212-2 (WIN, 5 μM) also induced a robust presynaptic depression of GABAergic inputs with no differences between genotypes (**Figure 3C**). These results indicate that the iLTD impairment in EAAT3-overexpressing mice arises from insufficient glutamate spillover to activate mGluRs, rather than a defect in mGluR expression, endocannabinoid signaling, or presynaptic depression mechanisms downstream of receptor activation.

**Figure 3.**
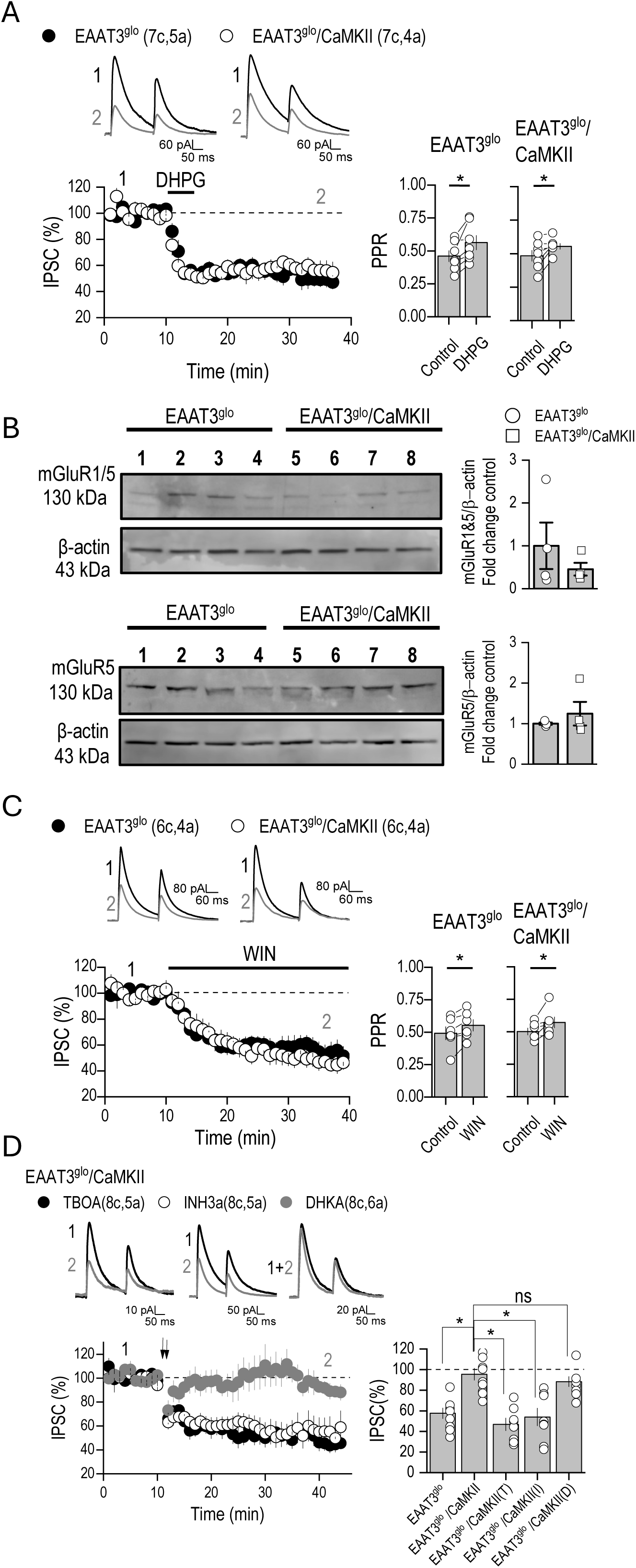
Blockade of EAAT3 restores impaired iLTD in EAAT3 overexpressing mice. **A,** Representative traces and summarized plots showing that bath application of group I mGluR agonist DHPG (5 min, 50 µM) depresses electrically evoked IPSCs to a similar extent in control (EAAT3^glo^) and EAAT3 overexpressing mice (EAAT3^glo^/CMKII). DHPG-induced depression of IPSCs is accompanied by changes in paired pulse ratio (PPR). **B**, Western blot analysis showing that the levels of group I mGluRs is similar between genotypes. **C**, Activation of presynaptic CB1R with WIN55,212-2 (WIN, 5 mM) suppresses GABAergic IPSC to a similar extent in both genotypes. WIN-mediated depression is accompanied by changes in PPR. **D**, DHKA (1 mM; D), a glial glutamate transporter inhibitor has no effect on the magnitude of iLTD, whereas TBOA (10 mM, T), a broad glutamate transporter inhibitor and the selective EAAT3 blocker INH3a (30 mM; I) rescue the iLTD impairments observed in slices of EAAT3 overexpressing mice. In panels A, C and D, data are presented as mean ± SEM, and representative traces taken at times indicated by numbers are shown. The number of cells (c) and animals (a) are indicated in parenthesis. *P <0.05.

Finally, we investigated the role of EAAT3 in altered heterosynaptic iLTD by evaluating the effect of various EAAT blockers on rescuing iLTD in overexpressing mice. The application of TBOA (10 μM), a broad glutamate transporter inhibitor, enhanced iLTD in control slices (**Supplementary Figure 3A**) and restored iLTD impairment in overexpressing slices (**Figure 3D**). In contrast, DHKA (1 μM), a glial glutamate transporter inhibitor had no effect on the magnitude of iLTD in EAAT3^glo^/CaMKII mice (**Figure 3D**) or in control mice (**Supplementary Figure 3A**). Interestingly, the selective EAAT3 blocker INH3a (30 μM) enhanced iLTD in control slices (**Supplementary Figure 3A**) and notably rescued the impairments observed in overexpressing slices (**Figure 3D**). Likewise, increased EAAT3 expression also impaired excitatory mGluR-mediated LTD, a phenomenon that could also be rescued by blocking EAAT3 with INH3a (**Supplementary Figure 3B**). Altogether, these results indicate that pharmacological blockade of EAAT3 (but not glial EAATs) is sufficient to normalize iLTD and further highlight that EAAT3 is the key transporter controlling the threshold for this form of heterosynaptic inhibitory plasticity in the hippocampus.

Given the roles of group I mGluRs and GABAergic plasticity in spatial memory and reversal learning (26, 33–35), we next evaluated whether the iLTD alterations observed in EAAT3^glo^/CaMKII mice were associated with changes in behavioral performance during spatial memory acquisition and cognitive flexibility in the Morris water maze (**Figure 4**). During the seven-day acquisition phase, no significant differences in the escape latency were observed between control and EAAT3^glo^/CaMKII mice treated with saline (**Figure 4B**). Consistently, pharmacological blockade of EAAT3 by systemic administration of INH3a did not affect performance during the acquisition phase in either control or EAAT3 overexpressing mice (**Figure 4C**). To quantitatively compare spatial navigation performance across training days, we analyzed the area under the curve (AUC) of the escape latency plots, which provides an integrated measure of spatial learning over time. During the acquisition phase, AUC values did not differ between genotypes or with INH3a treatment, indicating comparable overall performance. In contrast, during the reversal learning phase, EAAT3^glo^/CaMKII mice exhibited significantly longer escape latencies and higher AUC values compared to control mice (**Figure 4D**). Remarkably, EAAT3 blockade by INH3a rescued the reversal learning deficit observed in EAAT3 overexpressing mice, as evidenced by reduced escape latencies during the reversal phase (**Figure 4E**). On the spatial retention day, no significant differences were observed between control and EAAT3^glo^/CaMKII mice treated with either saline or INH3a during the probe trial of both the acquisition phase (**Supplementary Figure 5A**) and the reversal learning phase (**Supplementary Figure 5B**). Furthermore, and consistent with the lack of alterations in GABAergic synaptic plasticity when EAAT3 was overexpressed in GABAergic interneurons, no significant differences were observed between control and EAAT3^glo^/GAD mice during the acquisition (**Figure 4F**) or reversal (**Figure 4G**) phases of the spatial memory task. Together, these results indicate that EAAT3 overexpression in principal neurons is associated with a selective impairment in cognitive flexibility, while spatial learning acquisition remains unaffected, and show that this impairment can be rescued by selective EAAT3 inhibition.

**Figure 4.**
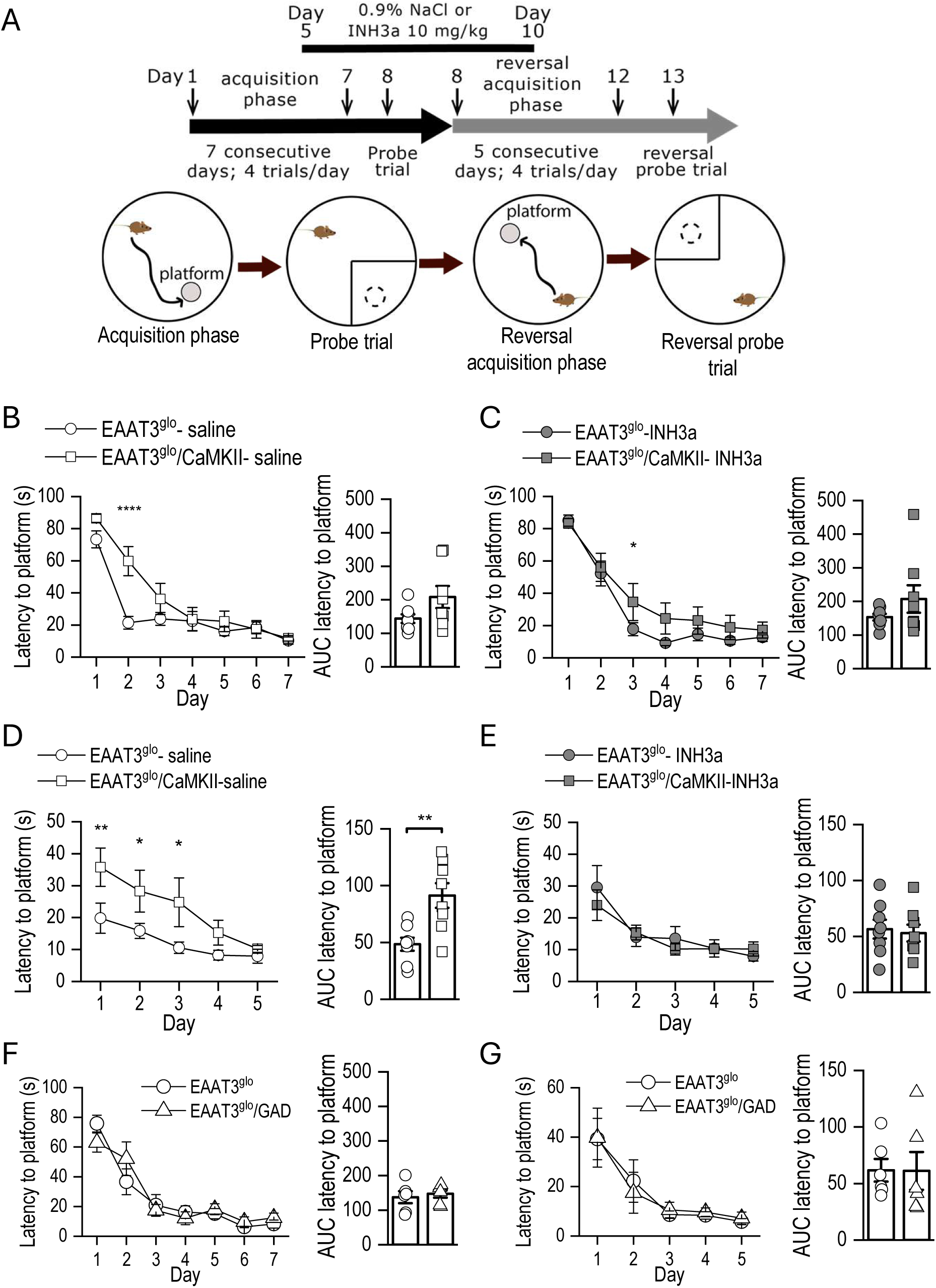
Increased expression of EAAT3 at principal neurons, but not GABAergic neurons impair hippocampal reversal learning. **A,** Diagram of experimental design of the Morris water maze acquisition and reversal learning paradigm. No differences were observed between (EAAT3^glo^) and EAAT3 overexpressing mice (EAAT3^glo^/CMKII) injected with saline **(B)** or EAAT3 blocker INH3a **(C)** in the time to reach the platform over the 7 days of the acquisition phase. **D**, saline-treated EAAT3^glo^/CaMKII mice exhibited significantly longer escape latencies compared with control mice, indicative of impaired reversal. **E**, the selective EAAT3 blocker INH3a rescues the impaired reversal in EAAT3 overexpressing mice. Data are expressed as mean ± SEM. Saline: EAAT3glo n = 8 and EAAT3glo/CaMKII n = 8; INH3a: EAAT3glo n = 8 and EAAT3glo/CaMKII n = 8. *P <0.05, **P <0.01 and ****P <0.0001.

## Discussion

Here we reveal that overexpression of the neuronal glutamate transporter EAAT3 in principal neurons, but not in GABAergic interneurons, controls the threshold of heterosynaptic inhibitory iLTD at hippocampal CA1 inhibitory synapses and produces a specific deficit in cognitive flexibility, without affecting basal excitatory or inhibitory transmission, excitatory/inhibitory balance, or spatial learning acquisition. Moreover, we found that pharmacological blockade of EAAT3, but not glial EAATs, the most abundant transporters expressed in forebrain and responsible for the majority of glutamate clearance (36), restores iLTD and rescues reversal learning performance in EAAT3- overexpressing mice. These findings identify EAAT3 as a regulator of glutamate spillover that sets the threshold for mGluR-dependent inhibitory plasticity. Consistent with this idea, overexpression of EAAT3 also controls the magnitude of mGluR-mediated LTD at homosynaptic excitatory synapses, a phenomenon that can be also rescued by pharmacological block of EAAT3. Thus, by controlling the threshold of synaptic LTD in the hippocampus, EAAT3 contributes to hippocampal-dependent reversal learning and establishes EAAT3 as a novel target for modulating hippocampal synaptic plasticity and adaptive behavior. Indeed, in humans, the correct activity of the hippocampus is necessary and required to respond in an appropriately flexible manner to high-order environments, and disruptions in this structure can render behavior habitual and inflexible (37).

A central mechanistic finding of our study is that EAAT3 overexpression in principal neurons abolishes heterosynaptic iLTD at CA1 GABAergic synapses, a form of synaptic plasticity that requires postsynaptic group I mGluR activation, endocannabinoid release, and presynaptic CB1R signaling(15). Importantly, several lines of evidence support the conclusion that this deficit arises from reduced glutamate spillovers rather than downstream signaling defects induced by EAAT3 overexpression. First, altered iLTD can be restored by exogenous application of glutamate when combined with a high electrical stimulation that, by itself, cannot induce iLTD **(Figure 2D)**, indicating that iLTD induction protocol fails because EAAT3 overexpression prevents sufficient glutamate accumulation and escape near mGluRs, but saturating transporters with exogenous glutamate can overcome this barrier. Second, direct activation of group I mGluRs or CB1Rs with exogenous application of agonists DHPG or WIN, respectively, induced robust presynaptic depression of GABAergic inputs with no genotype differences (**Figure 3A, C**). Moreover, group I mGluR expression levels were comparable between genotypes (**Figure 3B**), indicating that the mGluR–endocannabinoid–CB1R signaling cascade and presynaptic expression machinery are fully functional in EAAT3 overexpressing mice. Third, the selective EAAT3 inhibitor INH3a(38, 39) restored iLTD in overexpressing slices and increased the magnitude of iLTD in control mice (**Figure 3D**), whereas the glial EAAT1/2 blocker DHKA had no effect. In this regard, it has been shown that the non-selective glutamate transporter blocker TBOA, but not DHKA, alters excitatory NMDAR-dependent LTP in the hippocampus (40) supporting the idea that EAAT3-mediated uptake at excitatory synapses, rather than astrocytic clearance, might be a critical determinant of iLTD induction. Moreover, these findings are consistent with the idea that EAAT3 overexpression enhances glutamate clearance during high-frequency activity, thereby limiting the spatiotemporal spread of glutamate to extrasynaptic or perisynaptic group I mGluRs, limiting the probability of mGluR activation below the threshold required for iLTD induction. Accordingly, increased EAAT3 expression also controls the magnitude of mGluR-mediated LTD at homosynaptic excitatory synapses (**Supplementary Figure 3B**), highlighting the role of EAAT3 in regulating different forms of hippocampal LTD. Interestingly, increased GABAergic transmission in the hippocampus has been suggested to positively correlate with reversal learning(41), further supporting the idea that changes in inhibitory function might be important to specific aspects of hippocampal-dependent learning.

Behaviorally, EAAT3 overexpression produces a selective deficit in reversal learning in the Morris water maze, without impacting spatial memory acquisition or retention. This mirrors the electrophysiological phenotype: basal transmission and LTP-related processes (which support initial learning) are intact in EAAT3^glo^/CaMKII (42), whereas different forms of LTD required for updating or “unlearning” prior associations are impaired. This aligns with a growing body of evidence implicating LTD—both excitatory and inhibitory—in cognitive flexibility, reversal learning, and extinction (3–6, 8, 10). Notably, acute pharmacological blockade of EAAT3 with the selective inhibitor INH3a (38, 39) rescues the reversal learning deficit without affecting acquisition, demonstrating that inflexibility is directly caused by increased EAAT3 function and can be reversed in adulthood. These findings has potential translational relevance, as increased EAAT3 has been linked to OCD (23–25, 43), a neuropsychiatric disorder characterized by cognitive rigidity and impaired reversal learning (29, 44). Accordingly, other animal models relevant in OCD field including SAPAP3 knockout mice, also show impaired behavioral flexibility (27, 45), phenotypes that parallel the reversal learning deficits in EAAT3 overexpressing mice. Of note, SAPAP3 is a scaffolding protein involved in the regulation of postsynaptic mGluRs synaptic plasticity, through its effect on group I mGluR activity (46). Although alterations in mGluR-LTD have links to different neurological illnesses including mental retardation, autism, Alzheimer’s disease, Parkinson’s disease, and drug addiction (47), the parallels between in EAAT3 overexpressing and SAPAP3 knockout mice support the idea that disrupted mGluR-dependent plasticity in corticostriatal and hippocampal circuits could bea relevant pathophysiological mechanism in OCD, making both models valuable for studying synaptic bases of cognitive flexibility. Further experiments will be required to evaluate whether chronic EAAT3 inhibition might improve cognitive flexibility in disease models characterized by rigidity, including OCD.

## Methods

### Animals

Animal handling and use followed a protocol approved by the Animal Care and Use Committee of the Universidad de Valparaiso (BEA 219-25; CBC 127-2025) in accordance with the bioethics regulation of the Chilean Research Council (ANID).

### Hippocampal slice preparation

Acute transverse hippocampal slices (400 µm thick) were prepared from postnatal day 30 (P30) to P50, C57BL/6 mice control (EAAT3^glo^), EAAT3 overexpression at excitatory neurons (EAAT3^glo^/CMKII)(23) and inhibitory neurons (EAAT3^glo^/GAD) as previously described (48–50). EAAT3^glo^/GAD mice were generated by crossing EAAT3^glo^ mice with Gad2-IRES-Cre driver mice (Cat N° 028867; The Jackson Laboratory, Bar Harbor, ME, USA). In brief, the hippocampi were isolated from these animals and cut in a solution containing (in mM): 215 sucrose, 2.5 KCl, 26 NaHCO3, 1.6 NaH2PO4, 1 CaCl2, 4 MgCl2, 4 MgSO4 and 20 glucose. Thirty minutes post sectioning, the cutting medium was gradually switched to extracellular artificial cerebrospinal (ACSF) recording solution containing: 124 NaCl, 2.5 KCl, 26 NaHCO3, 1 NaH2PO4, 2.5 CaCl2, 1.3 MgSO4 and 10 glucose. All solutions were equilibrated with 95% O2 and 5% CO2 (pH 7.4). Slices were incubated for at least 30 min in the ACSF solution prior to recordings.

### Electrophysiology

All experiments, except where indicated, were performed at 28 ± 1°C in a submersion-type recording chamber perfused at ∼1-2 ml/min with ACSF supplemented with the AMPA and NMDA receptor antagonist CNQX and APV (25 μM), respectively. Whole-cell patch-clamp recordings using a Multiclamp 700A amplifier (Molecular Devices) were made from CA1 pyramidal neurons voltage clamped at 0 mV (unless otherwise stated) using patch-type pipette electrodes (∼3–4 MΩ) containing (in mM): 131 Cs-Gluconate, 8 NaCl, 1 CaCl2, 10 EGTA, 10 glucose, 10 HEPES; pH 7.2, 285-292 mmol/kg.

Miniature excitatory postsynaptic currents (mEPSCs) were recorded at 32 ± 1°C from neurons voltage-clamped at -60 mV in the continuous presence of tetrodotoxin (TTX, 500 nM). In contrast, isolated miniature inhibitory postsynaptic currents (mIPSCs) were recorded at 0 mV in the constant presence of TTX, CNQX (25 μM), and D-APV (50 μM) to block Na dependent release, AMPA and NMDA receptors, respectively. mEPSCs and mIPSC were identified using a minimal threshold amplitude (≥5 pA) and analyzed using the mini-analysis software Synaptosoft (Synaptosoft). Spontaneous excitatory and inhibitory currents (sEPSCs/sIPSCs) were recorded in absence of TTX, whereas AMPAR/NMDAR ratios were analyzed by recording AMPAR-mediated EPSCs at -60 mV and NMDAR-mediated EPSCs at +40mV in the continuous presence of CNQX (25 μM). To stimulate synaptic inputs, a monopolar stimulating patch-type pipettes were filled with ACSF and placed in the *stratium radiatum* (<50 μm from the CA1 pyramidal neurons).

Two different protocols were used to evaluate short-term synaptic plasticity: First, two pulses at different interstimulus intervals (10, 30, 70, 100, and 300 ms) were used to calculate the paired-pulse ratio (PPR) that was defined as the ratio of the amplitude of the second response to the amplitude of the first one. Second, synaptic depression was evaluated using a burst of 25 (for EPSC) and 20 (for IPSC) stimuli at 14 and 10 Hz, respectively, and delivered every 60 s; 5 burst-evoked responses were averaged for each experiment. Heterosynaptic iLTD was evoked using a standard high frequency stimulation consisting of a series of 100 stimuli (100 Hz) repeated 2 times at 20 sec intervals as previously described(15). HFS-iLTD was typically induced after 10 min of stable baseline and the magnitude of LTD was compared 25-30 min after LTD protocol. PPR was calculated 10 min before and 15 to 20 min after application of LTD induction protocol. In addition, paired-pulse low-frequency stimulation (900 paired-pulse, 50 ms interstimulus interval, at 1 Hz) was also used to test mGluR-mediated LTD at excitatory synapses. Drugs were obtained from Sigma, Tocris and Ascent Scientific, except INH3a that was acquired from MolPort SIA (Riga, LV-1011, Latvia). Stock solutions were added to the ACSF as needed. Total DMSO in the ACSF was maintained at <0.01% and 0.02%, respectively.

EPSCs and IPSCs were elicited at 20 s intervals, filtered at 2.2 kHz, and acquired at 5 kHz using a custom-made software written in Igor Pro XX (Wavemetrics, Inc., Lake Oswego, OR, USA). Series resistance (∼14-28 MΩ) was monitored throughout all experiments with a −5 mV, 80 ms voltage step, and cells that exhibited significant change in series resistance (>20%) were excluded from analysis. Statistical comparisons were made using unpaired and paired two-tailed Student’s t-test at the p<0.05 significance level in OriginPro 7.0 software (OriginLab Corporation, Northampton, MA). Unless otherwise indicated, all values are provided as the mean ± s.e.m. and illustrated traces are averages of 31-40 responses.

### Western blot analysis

We measured protein changes of mGluR1 and mGluR1/5 in hippocampal tissue using Western blot. In brief, mice were euthanized by rapid dislocation followed by decapitation to immediately remove the brains. Hippocampal samples were then homogenized in RIPA lysis buffer containing protease inhibitors (#78430, Thermo Fisher Scientific Inc, Waltham, MA, USA) and phosphatase inhibitors (J63907, Alfa Aesar, Tewksbury, MA, USA). The homogenates were incubated at 4°C for 30 min, sonicated and centrifuged at 14,000 × g at 4°C for 20 min, and the supernatants were collected. The concentration of proteins was measured by the Qubit™ Protein Assay Kit (Invitrogen). Crude protein extracts (50 μg) were denatured and separated on an 8% SDS-PAGE gel. After separation, the proteins were transferred on PVDF membranes (Amersham GE Healthcare, Bucks UK). The blots were blocked in 5% non-fat milk in phosphate-buffered saline with 0.05% Tween-20 for 1 hour at room temperature and incubated overnight at 4°C using antibody against mGluR1/5 (1:500; N75/33, NeuroMab), mGluR5 (1:500; AGC-007, Alomone), and β-actin (1:5000; ab8227, Abcam). These blots were incubated with appropriate horseradish peroxidase conjugated secondary antibodies and detected by ECL Select reagent (Amersham GE Healthcare, Bucks UK). The density of the selected bands was quantified using ImageJ software (National Institutes of Health, Bethesda, MD, USA). Statistical analysis was performed using unpaired Student’s t-test at the p<0.05 significance level in OriginPro 7.0 software (OriginLab Corporation, Northampton, MA). Data are expressed as mean ± s.e.m.

### Morris water maze

Morris water maze was performed as described previously(7). Mice were trained and evaluated in an indirectly illuminated room (20-40 lx) with salient cues located on the four walls. The maze consists of a circular pool (130 cm in diameter, 38 cm high) filled with water which was made opaque by addition of white nontoxic paint and maintained at 22–24 °C. A white, round plastic goal platform (15 cm in diameter) was submerged 0.5 cm below the water surface (28 cm high) in a fixed location in the NW, NE, SE, or SW quadrant 15 cm from the pool wall. Mice were trained in the acquisition phase for four trials per day with 20 min inter-trial intervals for 7 consecutive days. During each trial, mice were allowed to swim until they found the hidden platform, upon which they remained for 5 s before being returned to their home cage. Latency to reach the platform was recorded using Ethovision Video Tracking System (Noldus, The Netherlands). After the acquisition phase, the platform was removed from the pool, and mice were allowed to swim for 90 seconds in a single probe trial. Next, mice underwent reversal learning, in which the platform was moved to a new location within a different quadrant. Reversal learning continued for five consecutive days, following the same protocol as the acquisition phase. About 24 h after reversal learning completion, mice were evaluated to a second probe trial (reversal probe trial). During the probe trials, the time spent in the target quadrant (where the platform was located during acquisition or reversal learning) was recorded.

The selective EAAT3 blocker INH3a (38), dissolved in 0.9% NaCl and 1% DMSO (vehicle), was subcutaneously administered at a dose of 10 mg/kg one hour prior to the first trial between the fifth day of the acquisition phase and the third day of the reversal learning phase (6 consecutive days) in the Morris Water Maze test. On days 1 to 4 of the acquisition phase and on days 4 and 5 of the reversal learning phase, mice were administered the vehicle solution. Control groups received daily administration of the vehicle solution throughout both the acquisition and reversal learning phases. Statistical analysis was performed using GraphPad Prism 8 (GraphPad Software). Data are expressed as mean ± s.e.m. For comparisons involving multiple groups, Two-Way ANOVA, Two-Way Repeated Measures ANOVA, and Three-Way Repeated Measures ANOVA were used, followed by Tukey’s or uncorrected Fisher’s LSD post hoc analysis. A p<0.05 was considered statistically significant in all tests.

## Acknowledgments

This work was supported by the Chilean government through ANID FONDECYT Regular # 1252002 (A.E.C.), #1231012 (P.R.M.), Proyecto Puente UVA22991 (to A.E.C.) and ANID Millennium Institute CINV (ICN2025_026 to A.E.C). CA-G, N.A, S.F.E and W.P-B were supported by a PhD fellowship from ANID #21201603, # 21161701, 21191436 and #21220829, respectively.

## Author’s contributions

C.A-G, S.F.E, and A.A. performed all electrophysiological experiments and analyzed the results. N.M.A. and W.P-B. performed all behavioral experiments and analyzed the results. A.E.C. and P.R.M. designed experiments, guided the research, provided resources, and interpreted all the results. C.A-G., P.R.M. and A.E.C. wrote the paper.

## Supplementary Figure

**Supplementary Figure 1.** Excitatory synaptic function is normal in overexpressing EAAT3 mice. **A,** Representative traces and summarized plots showing that the amplitude and frequency of excitatory miniature activity (mEPSC) is normal in EAAT3 overexpressing mice (EAAT3^glo^/CMKII) compared to control littermate (EAAT3^glo^). **B**, Paired-pulse responses at 10, 30, 70, 100, and 300 ms interstimulus intervals (ISI) shows no differences between genotypes. **C**, Short-term synaptic responses evoked by a burst of 25 stimuli at 14 Hz is also similar between genotypes. **D**, No differences between genotypes on isolated NMDA input-output curve is observed. **E**, NMDAR-mediated EPSC show significant slower decay kinetics in EAAT3^glo^/CMKII mice compared to control littermate (EAAT3^glo^). **F,** EAAT3^glo^/CMKII synapses are more sensitive to the effect of GluN2B subunit antagonist Ro 25-6981 (5 μM) than control synapses. slices. In all panels, data are presented as mean ± SEM, and representative traces taken at times indicated by numbers are shown. Open circle in the bar graph represents a single cell. The number of cells (c) and animals (a) are indicated in parenthesis.

**Supplementary Figure 2.** Inhibitory synaptic function is normal in overexpressing EAAT3 mice. **A,** Representative traces and summarized plots showing that the amplitude and frequency of inhibitory miniature activity (mIPSC) is normal in EAAT3 overexpressing mice (EAAT3^glo^/CMKII) compared to control littermate (EAAT3^glo^). **B**, Paired-pulse responses at 10, 30, 70, 100, and 300 ms interstimulus intervals (ISI) shows no differences between genotypes. **C**, Synaptic responses and normalized summary data evoked by a burst of 20 stimuli at 10 Hz shows no difference between genotypes. **D**, GABA-mediated IPSC input-output curve is similar between genotypes Data are presented as mean ± SEM. The number of cells (c) and animals (a) are indicated in parenthesis.

**Supplementary Figure 3.** Inhibitory synaptic function is normal in mice overexpressing EAAT3 in GABAergic neurons. **A,** Representative traces and summarized plots showing that the amplitude and frequency of inhibitory spontaneous activity (sIPSC) is similar between EAAT3 overexpressing mice (EAAT3^glo^/GAD) and control littermate (EAAT3^glo^). **B**, However, an increase in the amplitude and frequency of miniature IPSC activity is observed between genotypes. *P <0.05. **C**, Evoked IPSC amplitudes as a function of stimulus intensity plotted as input/output curves in inhibitory synapses shows no differences between genotypes. **D**, Paired-pulse responses at 10, 30, 70, 100, and 300 ms interstimulus intervals (ISI) shows no differences between genotypes. **E**, DHPG (50 μM, 5 min)-induced a similar depression of IPSC between genotypes. Representative traces taken at times indicated by numbers are shown Data are presented as mean ± SEM. The number of cells (c) and animals (a) are indicated in parenthesis.

**Supplementary Figure 4.** Blocking EAAT3 increased the magnitude of iLTD and restored mGluR-dependent LTD at excitatory synapses in the CA1 area of the hippocampus. **A,** Representative traces (top) and summarized plots (bottom) showing that bath application of TBOA (10 mM), a broad glutamate transporter inhibitor and the selective EAAT3 blocker INH3a (30 mM) enhance the magnitude of iLTD elicited by high frequency stimulation (arrows), whereas, DHKA (1 mM), a glial glutamate transporter inhibitor had no effect on the magnitude of iLTD in control mice. **B**, Paired-pulse low frequency stimulation (PP-LFS; 900 paired pulse, 50 ms interval, at 1 Hz) elicited a robust mGluR-LTD at excitatory Schaffer collateral-to-CA1 synapses in the hippocampus. Excitatory mGluR-LTD is diminished in EAAT3 overexpressing mice (EAAT3^glo^/CaMKII) compared to control littermate (EAAT3^glo^) and can be restored to a similar extent by blocking EAAT3 with the selective blocker INH3a (30 mM). Data are presented as mean ± SEM, and representative traces taken at times indicated by numbers are shown. Each circle in the bar graph represents a single experiment. The number of cells (c) and animals (a) are indicated in parenthesis. *P <0.05.

**Supplementary Figure 5.** Systemic administration of EAAT3 blocker INH3a does not alter spatial retention in the probe trial of acquisition and reversal learning phases of the Morris water maze. **A,** No significant differences were observed in EAAT3^glo^ and EAAT3^glo^/CaMKII mice treated with EAAT3 blocker in the exploration time in the target quadrant on the probe trial day of the acquisition phase (A) or during the probe trial of the reversal learning phase. **B**, Data are expressed as mean ± SEM. The statistical analysis used was a Two-Way ANOVA followed by Tukey multiple comparisons analysis. EAAT3^glo^ - saline n = 8, EAAT3^glo^/CaMKII - saline n = 8, EAAT3^glo^ – INH3a n = 8 and EAAT3^glo^/CaMKII – INH3a n = 8.

## References

1. J. C. Magee, C. Grienberger, Synaptic Plasticity Forms and Functions. Annu Rev Neurosci 43, 95–117 (2020).

2. T. V. Bliss, G. L. Collingridge, A synaptic model of memory: long-term potentiation in the hippocampus. Nature 361, 31–39 (1993).

3. K. L. Eales et al., The MK2/3 cascade regulates AMPAR trafficking and cognitive flexibility. Nat Commun 5, 4701 (2014).

4. G. L. Collingridge, S. Peineau, J. G. Howland, Y. T. Wang, Long-term depression in the CNS. Nat Rev Neurosci 11, 459–473 (2010).

5. S. Duffy, V. Labrie, J. C. Roder, D-serine augments NMDA-NR2B receptor-dependent hippocampal long-term depression and spatial reversal learning. Neuropsychopharmacology 33, 1004–1018 (2008).

6. E. Morice et al., Parallel loss of hippocampal LTD and cognitive flexibility in a genetic model of hyperdopaminergia. Neuropsychopharmacology 32, 2108–2116 (2007).

7. T. Tsetsenis et al., Rab3B protein is required for long-term depression of hippocampal inhibitory synapses and for normal reversal learning. Proc Natl Acad Sci U S A 108, 14300–14305 (2011).

8. Z. Dong et al., Hippocampal long-term depression mediates spatial reversal learning in the Morris water maze. Neuropharmacology 64, 65–73 (2013).

9. D. Manahan-Vaughan, K. H. Braunewell, Novelty acquisition is associated with induction of hippocampal long-term depression. Proc Natl Acad Sci U S A 96, 8739–8744 (1999).

10. F. Mills et al., Cognitive flexibility and long-term depression (LTD) are impaired following beta-catenin stabilization in vivo. Proc Natl Acad Sci U S A 111, 8631–8636 (2014).

11. R. E. Nicholls et al., Transgenic mice lacking NMDAR-dependent LTD exhibit deficits in behavioral flexibility. Neuron 58, 104–117 (2008).

12. A. Awasthi et al., Synaptotagmin-3 drives AMPA receptor endocytosis, depression of synapse strength, and forgetting. Science 363 (2019).

13. K. E. Ireton et al., Regulation of the Ca(2+) Channel Ca(V)1.2 Supports Spatial Memory and Its Flexibility and LTD. J Neurosci 43, 5559–5573 (2023).

14. S. H. Oliet, R. C. Malenka, R. A. Nicoll, Two distinct forms of long-term depression coexist in CA1 hippocampal pyramidal cells. Neuron 18, 969–982 (1997).

15. V. Chevaleyre, P. E. Castillo, Heterosynaptic LTD of hippocampal GABAergic synapses: a novel role of endocannabinoids in regulating excitability. Neuron 38, 461–472 (2003).

16. P. E. Castillo, T. J. Younts, A. E. Chavez, Y. Hashimotodani, Endocannabinoid signaling and synaptic function. Neuron 76, 70–81 (2012).

17. S. Valtcheva, L. Venance, Control of Long-Term Plasticity by Glutamate Transporters. Front Synaptic Neurosci 11, 10 (2019).

18. S. Holmseth et al., The density of EAAC1 (EAAT3) glutamate transporters expressed by neurons in the mammalian CNS. J Neurosci 32, 6000–6013 (2012).

19. J. D. Rothstein et al., Localization of neuronal and glial glutamate transporters. Neuron 13, 713–725 (1994).

20. S. Bellini et al., Neuronal Glutamate Transporters Control Dopaminergic Signaling and Compulsive Behaviors. J Neurosci 38, 937–961 (2018).

21. A. Scimemi, H. Tian, J. S. Diamond, Neuronal transporters regulate glutamate clearance, NMDA receptor activation, and synaptic plasticity in the hippocampus. J Neurosci 29, 14581–14595 (2009).

22. J. R. Wendland et al., A haplotype containing quantitative trait loci for SLC1A1 gene expression and its association with obsessive-compulsive disorder. Arch Gen Psychiatry 66, 408–416 (2009).

23. C. Delgado-Acevedo et al., Behavioral and synaptic alterations relevant to obsessive-compulsive disorder in mice with increased EAAT3 expression. Neuropsychopharmacology 44, 1163–1173 (2019).

24. M. O. Chohan et al., Developmental impact of glutamate transporter overexpression on dopaminergic neuron activity and stereotypic behavior. Mol Psychiatry 27, 1515–1526 (2022).

25. J. M. Kopelman et al., Forebrain EAAT3 Overexpression Increases Susceptibility to Amphetamine-Induced Repetitive Behaviors. eNeuro 11 (2024).

26. Z. Yang et al., Dysfunction of Orbitofrontal GABAergic Interneurons Leads to Impaired Reversal Learning in a Mouse Model of Obsessive-Compulsive Disorder. Curr Biol 31, 381–393 e384 (2021).

27. E. E. Manning, A. Y. Dombrovski, M. M. Torregrossa, S. E. Ahmari, Impaired instrumental reversal learning is associated with increased medial prefrontal cortex activity in Sapap3 knockout mouse model of compulsive behavior. Neuropsychopharmacology 44, 1494–1504 (2019).

28. B. J. G. van den Boom, A. H. Mooij, I. Miseviciute, D. Denys, I. Willuhn, Behavioral flexibility in a mouse model for obsessive-compulsive disorder: Impaired Pavlovian reversal learning in SAPAP3 mutants. Genes Brain Behav 18, e12557 (2019).

29. P. Gruner, C. Pittenger, Cognitive inflexibility in Obsessive-Compulsive Disorder. Neuroscience 345, 243–255 (2017).

30. J. Gottwald et al., Impaired cognitive plasticity and goal-directed control in adolescent obsessive-compulsive disorder. Psychol Med 48, 1900–1908 (2018).

31. C. Pittenger, Biological Mechanisms and Treatment of Obsessive-Compulsive Disorder. Annu Rev Clin Psychol 22, 455–480 (2026).

32. G. C. Mathews, J. S. Diamond, Neuronal glutamate uptake Contributes to GABA synthesis and inhibitory synaptic strength. J Neurosci 23, 2040–2048 (2003).

33. J. Xu, Y. Zhu, A. Contractor, S. F. Heinemann, mGluR5 has a critical role in inhibitory learning. J Neurosci 29, 3676–3684 (2009).

34. G. Wiera et al., Long-term plasticity of inhibitory synapses in the hippocampus and spatial learning depends on matrix metalloproteinase 3. Cell Mol Life Sci 78, 2279–2298 (2021).

35. D. Manahan-Vaughan, K. H. Braunewell, The metabotropic glutamate receptor, mGluR5, is a key determinant of good and bad spatial learning performance and hippocampal synaptic plasticity. Cereb Cortex 15, 1703–1713 (2005).

36. R. J. Vandenberg, R. M. Ryan, Mechanisms of glutamate transport. Physiol Rev 93, 1621–1657 (2013).

37. A. Vila-Ballo et al., Unraveling the Role of the Hippocampus in Reversal Learning. J Neurosci 37, 6686–6697 (2017).

38. P. Wu, W. E. Bjorn-Yoshimoto, M. Staudt, A. A. Jensen, L. Bunch, Identification and Structure-Activity Relationship Study of Imidazo[1,2-a]pyridine-3-amines as First Selective Inhibitors of Excitatory Amino Acid Transporter Subtype 3 (EAAT3). ACS Chem Neurosci 10, 4414–4429 (2019).

39. L. van Veggel et al., EAAT3 modulation: A potential novel avenue towards remyelination in multiple sclerosis. Biomed Pharmacother 186, 117960 (2025).

40. J. R. Barnes et al., The Relationship Between Glutamate Dynamics and Activity-Dependent Synaptic Plasticity. J Neurosci 40, 2793–2807 (2020).

41. F. Morellini et al., Improved reversal learning and working memory and enhanced reactivity to novelty in mice with enhanced GABAergic innervation in the dentate gyrus. Cereb Cortex 20, 2712–2727 (2010).

42. N. M. Ardiles et al., Increased forebrain EAAT3 expression confers resilience to chronic stress. J Neurochem 169, e16216 (2025).

43. A. P. Escobar, J. R. Wendland, A. E. Chavez, P. R. Moya, The Neuronal Glutamate Transporter EAAT3 in Obsessive-Compulsive Disorder. Front Pharmacol 10, 1362 (2019).

44. G. Valerius, A. Lumpp, A. K. Kuelz, T. Freyer, U. Voderholzer, Reversal learning as a neuropsychological indicator for the neuropathology of obsessive compulsive disorder? A behavioral study. J Neuropsychiatry Clin Neurosci 20, 210–218 (2008).

45. J. M. Welch et al., Cortico-striatal synaptic defects and OCD-like behaviours in Sapap3-mutant mice. Nature 448, 894–900 (2007).

46. M. Chen et al., Sapap3 deletion anomalously activates short-term endocannabinoid-mediated synaptic plasticity. J Neurosci 31, 9563–9573 (2011).

47. C. Luscher, K. M. Huber, Group 1 mGluR-dependent synaptic long-term depression: mechanisms and implications for circuitry and disease. Neuron 65, 445–459 (2010).

48. C. Ancaten-Gonzalez et al., BK channels mediate a presynaptic form of mGluR-LTD in the neonatal hippocampus. Proc Natl Acad Sci U S A 122, e2411506122 (2025).

49. B. Brauer et al., Impact of KDM6B mosaic brain knockout on synaptic function and behavior. Sci Rep 14, 20416 (2024).

50. A. E. Chavez, V. M. Hernandez, A. Rodenas-Ruano, C. S. Chan, P. E. Castillo, Compartment-specific modulation of GABAergic synaptic transmission by TRPV1 channels in the dentate gyrus. J Neurosci 34, 16621–16629 (2014).

