## Supplementary figures and images for "Neuronal glutamate transporter EAAT3 regulates hippocampal GABAergic plasticity and reversal learning"

### Supplementary Figure 1

#### Ancaten et al., Supplementary Figure 1

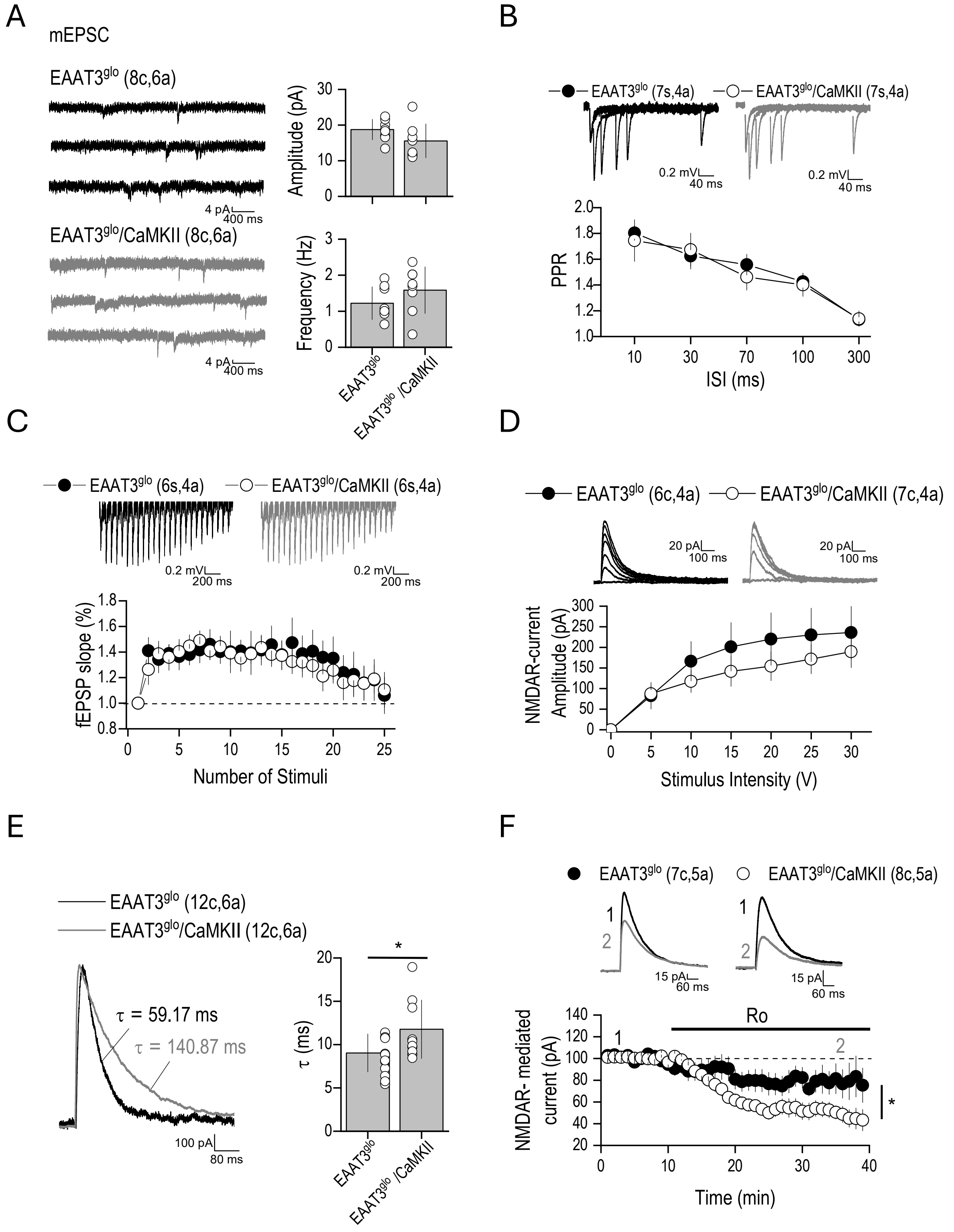

### Supplementary Figure 2

#### Ancaten et al., Supplementary Figure 2

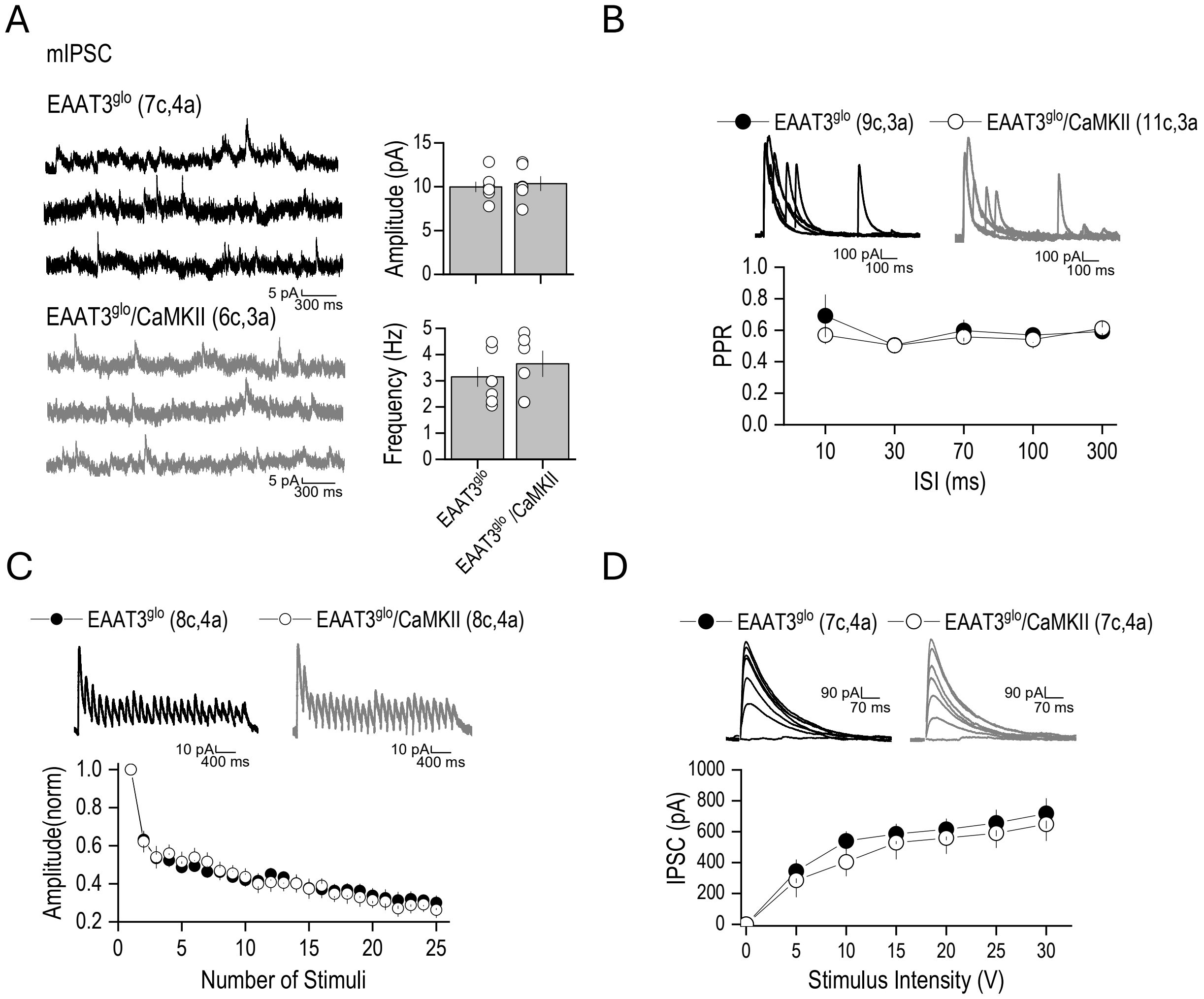

### Supplementary Figure 3

#### Ancaten et al., Supplementary Figure 3

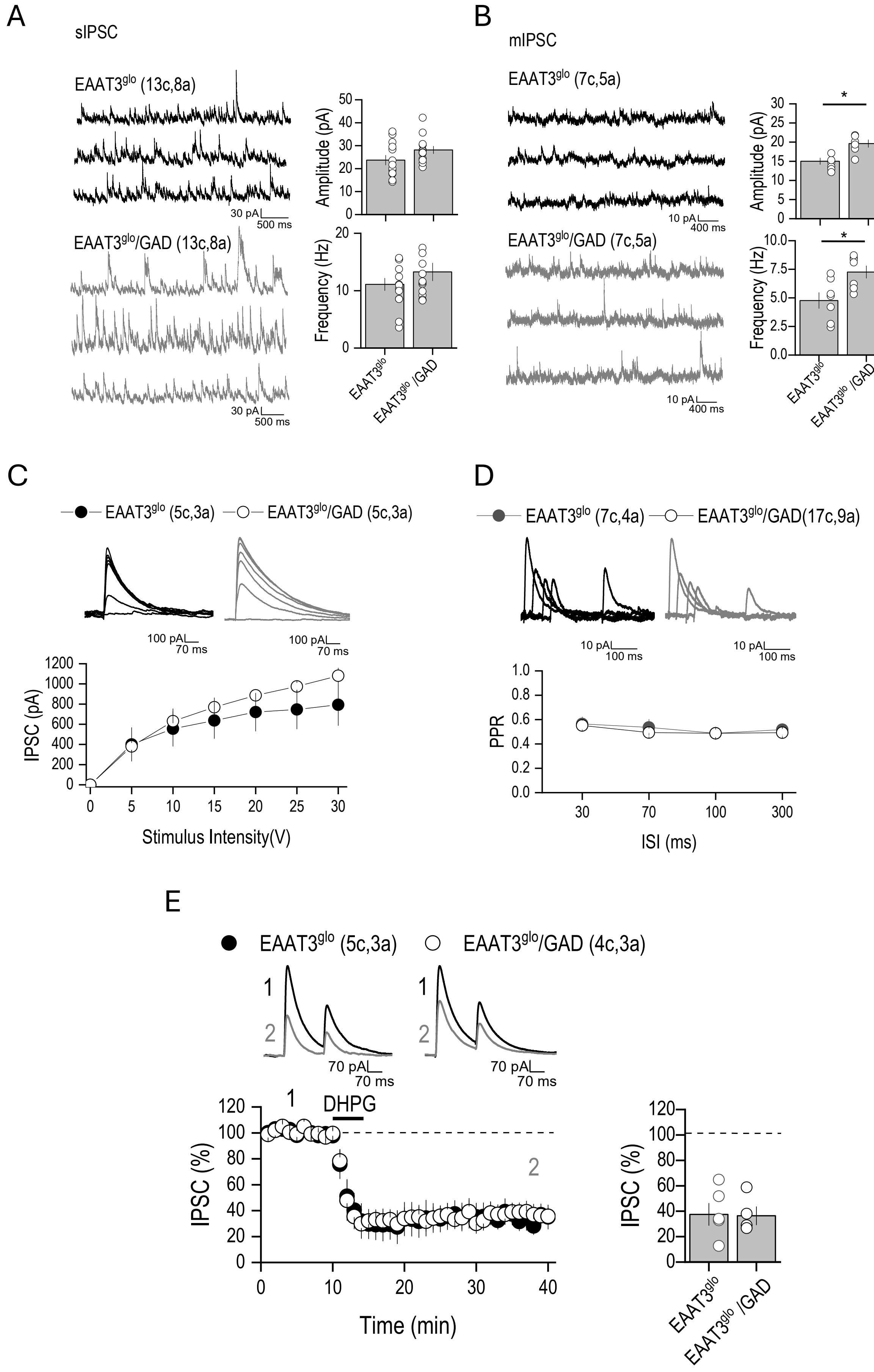

### Supplementary Figure 4

Ancaten et al., Supplementary Figure 4

A

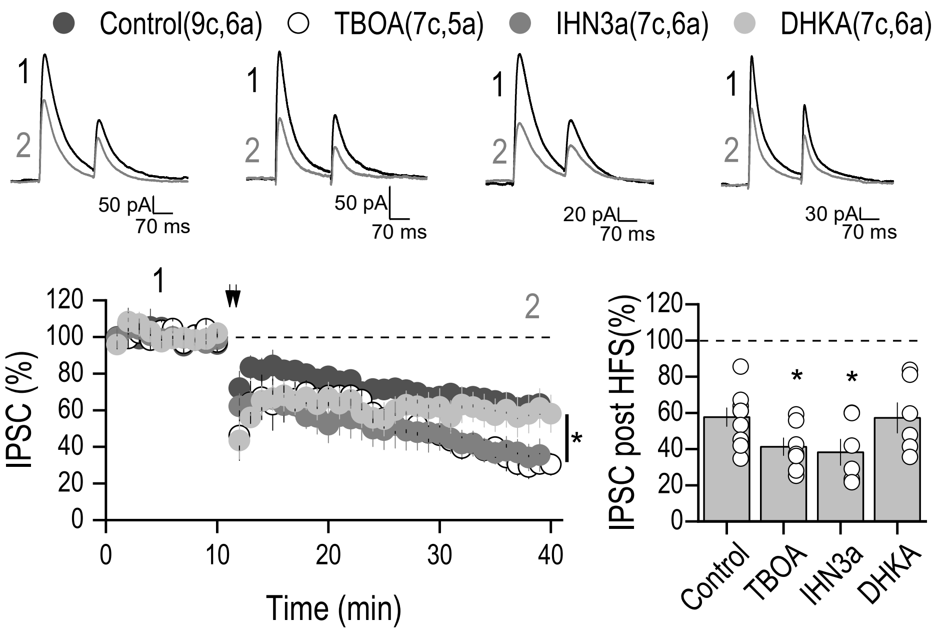


B

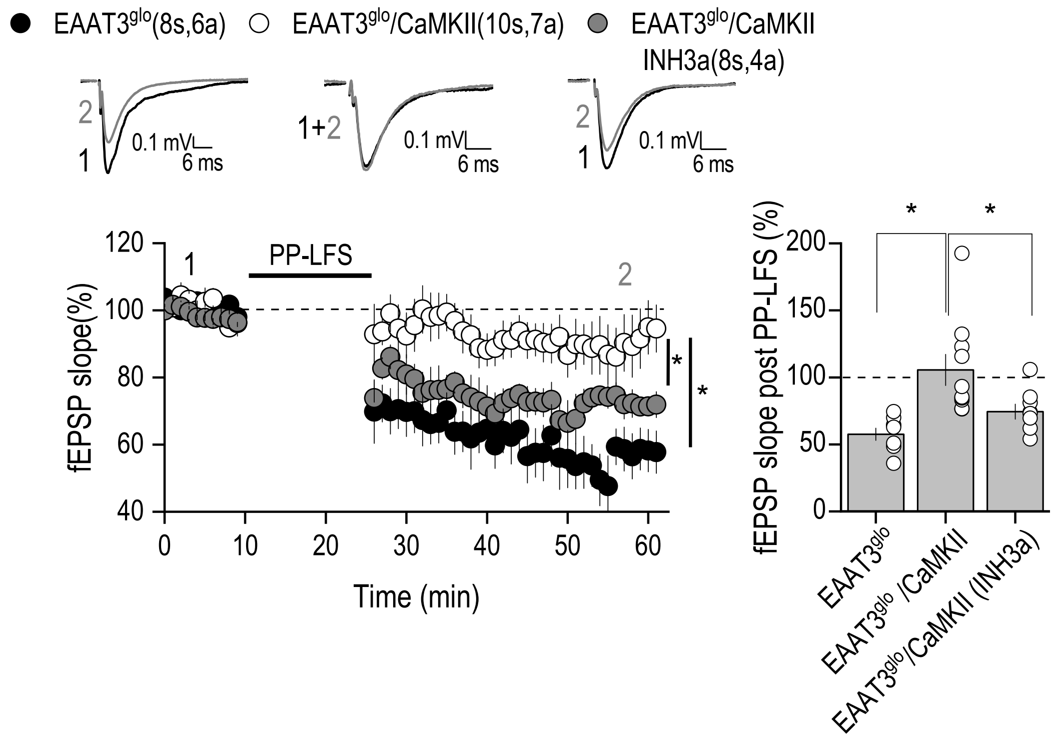

### Supplementary Figure 5

Ancaten et al., Supplementary Figure 5

A

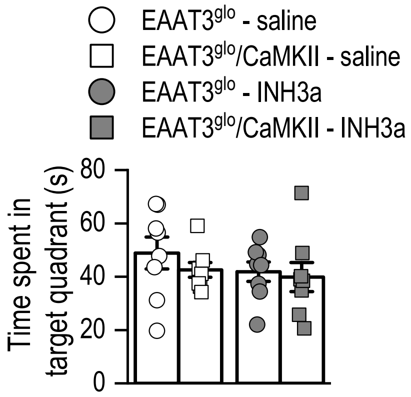


B

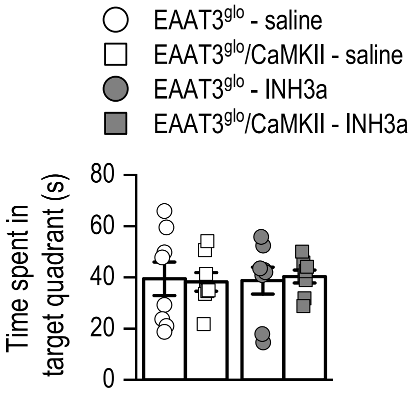
