## Supplementary Table 1 and 2 for "Neuronal glutamate transporter EAAT3 regulates hippocampal GABAergic plasticity and reversal learning"

**Supplementary Table 1**: **Quantitative analysis of cellular mechanism involved in mGluR-mediated LTD at GABAergic synapses in the hippocampus.** The table includes quantitative analyses of slice electrophysiology experiments shown in Figs. 1, 2 and 3. "n" represents number of slices (s), cells (c) and animals (a). PPR, paired-pulse ratio; Glu, Glutamate; HFS, high frequency stimulation; DHPG, group I mGluR agonist; WIN, WIN55,212-2 a CB1R agonist.

| Parameter | Mouse line | Value<br>(mean ± s.e.m) | N | P value | Figure |
|---|---|---|---|---|---|
| sEPSC amplitude | EAAT3$^{glo}$<br>EAAT3$^{glo}$/CaMKII | 16.84 ± 1.20<br>15.01 ± 1.01 | 6c/4a<br>8c/5a | P = 0.069 | 1A |
| sEPSC frequency | EAAT3$^{glo}$<br>EAAT3$^{glo}$/CaMKII | 5.09 ± 0.87<br>4.39 ± 0.62 | 6c/4a<br>8c/5a | P = 0.512 | 1A |
| AMPA/NMDA ratio | EAAT3$^{glo}$<br>EAAT3$^{glo}$/CaMKII | 0.96 ± 0.21<br>1.03 ± 0.23 | 17c/8a<br>15/8a | P = 0.801 | 1B |
| sIPSC amplitude | EAAT3$^{glo}$<br>EAAT3$^{glo}$/CaMKII | 21.69 ± 3.04<br>23.19 ± 2.83 | 7c/5a<br>7c/5a | P = 0.723 | 1C |
| sIPSC frequency | EAAT3$^{glo}$<br>EAAT3$^{glo}$/CaMKII | 13.44 ± 2.07<br>15.35 ± 2.46 | 7c/5a<br>7c/5a | P = 0.563 | 1C |
| Exciatroy/Inhibitory Balance | EAAT3$^{glo}$<br>EAAT3$^{glo}$/CaMKII | 0.45 ± 0.13<br>0.55 ± 0.17 | 6c/4a<br>6c/4a | P = 0.664 | 1C |
| Maginutde of GABAergic iLTD | EAAT3$^{glo}$<br>EAAT3$^{glo}$/CaMKII | 57.63 ± 5.08<br>95.53 ± 5.72 | 9c/6a<br>9c/6a | P < 0.001 | 2A |
| iLTD PPR | EAAT3$^{glo}$<br><br><br>EAAT3$^{glo}$/CaMKII | Pre 0.42 ± 0.05<br>Post 0.49 ± 0.03<br><br>Pre 0.56 ± 0.07<br>Post 0.55 ± 0.06 | 9c/6a<br>9c/6a<br><br>9c/6a<br>9c/6a | P = 0.024<br><br><br>P = 0.851 | 2A |
| Maginutde of GABAergic iLTD | EAAT3$^{glo}$<br>EAAT3$^{glo}$/GAD | 50.95 ± 3.80<br>48.50 ± 9.33 | 8c/5a<br>8c/5a | P =0.811 | 2B |
| iLTD PPR | EAAT3$^{glo}$<br><br><br>EAAT3$^{glo}$/GAD | Pre 0.48 ± 0.04<br>Post 0.59 ± 0.06<br><br>Pre 0.56 ± 0.07<br>Post 0.55 ± 0.06 | 8c/5a<br>8c/5a<br><br>8c/5a<br>8c/5a | P = 0.019<br><br><br>P < 0.001 | 2B |
| Glut (50 $\mu$M)<br>Glut + 1 HFS train | EAAT3$^{glo}$ | Glut 92.52 ± 3.68<br>Glut+HFS 30.20± 5.30 | 5c/4a<br>4c/3a | P < 0.001 | 2C |
| Glut (50 $\mu$M)<br>Glut + 1 HFS train<br>PPR | EAAT3$^{glo}$ | Glut Pre 0.50 ± 0.04<br>Glut Post 0.52 ± 0.01<br><br>Glu+HFS<br>Pre 0.54 ±0.03<br>Post 0.68 ± 0.05 | 5c/4a<br><br><br>4c/3a | P = 0.561<br><br><br>P = 0.040 | 2C |
| Glut (50 $\mu$M)<br>Glut + 1 HFS train | EAAT3$^{glo}$/CaMKII | Glut 105.35 ± 13.62<br>Glut+HFS 53.32± 15.09 | 5c/3a<br>5c/3a | P = 0.031 | 2D |
| Glut (50 $\mu$M)<br>Glut + 1 HFS train<br>PPR | EAAT3$^{glo}$/CaMKII | Glut Pre 0.53 ± 0.04<br>Glut Post 0.53 ± 0.02<br><br>Glu+HFS<br>Pre 0.46 ± 0.04<br>Post 0.59 ± 0.02 | 5c/3a<br><br><br>5c/3a | P = 0.883<br><br><br>P = 0.030 | 2D |

| | | | | | |
|---|---|---|---|---|---|
| DHPG-induced depression of IPSCs | EAAT3$^{glo}$<br>EAAT3$^{glo}$/CaMKII | 44.83 ± 4.84<br>54.16 ± 6.54 | 7c/5a<br>7c/4a | P = 0.284 | 3A |
| DHPG-induced depression of IPSCs<br>PPR | EAAT3$^{glo}$<br><br><br>EAAT3$^{glo}$/CaMKII | Pre 0.46 ± 0.04<br>Post 0.563 ± 0.05<br><br>Pre 0.49 ± 0.03<br>Post 0.55 ± 0.02 | 7c/5a<br><br><br>7c/4a | P = 0.013<br><br><br>P = 0.030 | 3A |
| WIN-induced depression of IPSCs | EAAT3$^{glo}$<br>EAAT3$^{glo}$/CaMKII | 55.87 ± 7.19<br>48.01 ± 4.79 | 6c/4a<br>6c/4a | P = 0.384 | 3C |
| WIN-induced depression of IPSCs<br>PPR | EAAT3$^{glo}$<br><br><br>EAAT3$^{glo}$/CaMKII | Pre 0.49 ± 0.05<br>Post 0.55 ± 0.04<br><br>Pre 0.50 ± 0.02<br>Post 0.57 ± 0.04 | 6c/4a<br><br><br>6c/4a | P = 0.013<br><br><br>P = 0.040 | 3C |
| Glutamate transporter blockers on iLTD magnitude | EAAT3$^{glo}$/CaMKII | TBOA 46.85 ± 6.04<br>DHKA 88.33 ± 4.84<br>INH3 54.03 ± 8.50<br><br>TBOA vs DHKA<br>TBOA vs INH3<br>DHKA vs INH3 | 8c/6a<br>8c/5a<br>8c/6a | P = 0.020<br>P = 0.449<br>P = 0.020<br><br>P < 0.001<br>P = 0.503<br>P = 0.007 | 3D |

**Supplementary Table 2: Quantitative analysis of cellular mechanisms involved in mGluR-mediated LTD at GABAergic synapses in the hippocampus**. The table includes quantitative analyses of experiments shown in Supplementary Figures 1 to 4. “n” represents number of cells (c), slices (s) and animals (a). PPR, paired-pulse ratio;

| Parameter | Mouse line | Value (mean ± s.e.m) | N | P value | Figure |
|---|---|---|---|---|---|
| mEPSC amplitude | $EAAT3^{glo}$<br>$EAAT3^{glo}$/CaMKII | 18.76 ± 2.87<br>15.56 ± 4.75 | 8c/6a<br>8c/6a | P = 0.145 | Sup. 1A |
| mEPSC frequency | $EAAT3^{glo}$<br>$EAAT3^{glo}$/CaMKII | 1.22 ± 0.45<br>1.59 ± 0.64 | 8c/6a<br>8c/6a | P = 0.278 | Sup. 1A |
| fEPSP<br>PPR 10 ms | $EAAT3^{glo}$<br>$EAAT3^{glo}$/CaMKII | 1.80 ± 0.05<br>1.74 ± 0.16 | 7s/4a<br>7s/4a | P = 0.760 | Sup. 1B |
| fEPSP<br>PPR 30 ms | $EAAT3^{glo}$<br>$EAAT3^{glo}$/CaMKII | 1.63 ± 0.10<br>1.67 ± 0.20 | 7s/4a<br>7s/4a | P = 0.788 | Sup. 1B |
| fEPSP<br>PPR 70 ms | $EAAT3^{glo}$<br>$EAAT3^{glo}$/CaMKII | 1.55 ± 0.08<br>1.46 ± 0.10 | 7s/4a<br>7s/4a | P = 0.546 | Sup. 1B |
| fEPSP<br>PPR 100 ms | $EAAT3^{glo}$<br>$EAAT3^{glo}$/CaMKII | 1.43 ± 0.06<br>1.40 ± 0.09 | 7s/4a<br>7s/4a | P = 0.850 | Sup. 1B |
| fEPSP<br>PPR 300 ms | $EAAT3^{glo}$<br>$EAAT3^{glo}$/CaMKII | 1.13 ± 0.01<br>1.13 ± 0.06 | 7s/4a<br>7s/4a | P = 0.948 | Sup. 1B |
| fEPSP Train ratio 25th /1s | $EAAT3^{glo}$<br>$EAAT3^{glo}$/CaMKII | 1.06 ± 0.14<br>1.10 ± 0.14 | 6s/4a<br>6s/4a | P = 0.668 | Sup. 1C |
| NMDA current input/output curve 5 V | $EAAT3^{glo}$<br>$EAAT3^{glo}$/CaMKII | 82.77 ± 32.46<br>87.41 ± 18.65 | 6c/4a<br>7c/4a | P = 0.934 | Sup. 1D |
| NMDA current input/output curve 10 V | $EAAT3^{glo}$<br>$EAAT3^{glo}$/CaMKII | 166.52 ± 47.91<br>117.55 ± 27.12 | 6c/4a<br>7c/4a | P = 0.256 | Sup. 1D |
| NMDA current input/output curve 15 V | $EAAT3^{glo}$<br>$EAAT3^{glo}$/CaMKII | 201.46 ± 58.95<br>141.65 ± 36.38 | 6c/4a<br>7c/4a | P = 0.213 | Sup. 1D |
| NMDA current input/output curve 20 V | $EAAT3^{glo}$<br>$EAAT3^{glo}$/CaMKII | 220.17 ± 63.84<br>154.49 ± 35.20 | 6c/4a<br>7c/4a | P = 0.186 | Sup. 1D |
| NMDA current input/output curve 25 V | $EAAT3^{glo}$<br>$EAAT3^{glo}$/CaMKII | 230.25 ± 64.83<br>171.54 ± 35.71 | 6c/4a<br>7c/4a | P = 0.297 | Sup. 1D |
| NMDA current input/output curve 30 V | $EAAT3^{glo}$<br>$EAAT3^{glo}$/CaMKII | 171.54 ± 35.71<br>189.58 ± 38.82 | 6c/4a<br>7c/4a | P = 0.452 | Sup. 1D |
| NMDA receptor kinetics | $EAAT3^{glo}$<br>$EAAT3^{glo}$/CaMKII | 9.03 ± 2.18<br>11.77 ± 3.37 | 12c/6a<br>12c/6a | P = 0.028 | Sup. 1E |
| Ro 25-6981 effect on NMDA receptor amplitude | $EAAT3^{glo}$<br>$EAAT3^{glo}$/CaMKII | 78.82 ± 9.30<br>47.97 ± 6.98 | 7c/5a<br>8c/5a | P = 0.017 | Sup. 1F |
| mIPSC amplitude | $EAAT3^{glo}$ | 9.98 ± 0.57 | 7c/4a | P = 0.709 | Sup. 2A |

| | | | | | |
|---|---|---|---|---|---|
| | EAAT3glo/CaMKII | 10.35 ± 0.83 | 6c/3a | | |
| mIPSC frequency | EAAT3glo<br>EAAT3glo/CaMKII | 3.15 ± 0.37<br>3.64 ± 0.49 | 7c/4a<br>6c/3a | P = 0.429 | Sup. 2A |
| IPSCs<br>PPR 10 ms | EAAT3glo<br>EAAT3glo/CaMKII | 0.69 ± 0.13<br>0.57 ± 0.05 | 9c/3a<br>11c/3a | P = 0.374 | Sup. 2B |
| IPSCs<br>PPR 30 ms | EAAT3glo<br>EAAT3glo/CaMKII | 0.51 ± 0.04<br>0.50 ± 0.04 | 9c/3a<br>11c/3a | P = 0.974 | Sup. 2B |
| IPSCs<br>PPR 70 ms | EAAT3glo<br>EAAT3glo/CaMKII | 0.60 ± 0.06<br>0.56 ± 0.04 | 9c/3a<br>11c/3a | P = 0.603 | Sup. 2B |
| IPSCs<br>PPR 100 ms | EAAT3glo<br>EAAT3glo/CaMKII | 0.57 ± 0.04<br>0.54 ± 0.05 | 9c/3a<br>11c/3a | P = 0.715 | Sup. 2B |
| IPSCs<br>PPR 300 ms | EAAT3glo<br>EAAT3glo/CaMKII | 0.59 ± 0.02<br>0.61 ± 0.03 | 9c/3a<br>11c/3a | P = 0.654 | Sup. 2B |
| IPSC Train ratio<br>25th /1s | EAAT3glo<br>EAAT3glo/CaMKII | 0.30 ± 0.04<br>0.26 ± 0.04 | 8c/4a<br>8c/4a | P = 0.500 | Sup.2C |
| IPSC input/output<br>curve<br>5 V | EAAT3glo<br>EAAT3glo/CaMKII | 344.00 ± 74.72<br>285.48 ± 108.35 | 7c/4a<br>7c/4a | P = 0.673 | Sup 2D |
| IPSC input/output<br>curve<br>10 V | EAAT3glo<br>EAAT3glo/CaMKII | 539.01 ± 62.12<br>403.50 ± 91.10 | 7c/4a<br>7c/4a | P = 0.254 | Sup 2D |
| IPSC input/output<br>curve<br>15 V | EAAT3glo<br>EAAT3glo/CaMKII | 585.58 ± 64.47<br>528.53 ± 106.49 | 7c/4a<br>7c/4a | P = 0.665 | Sup 2D |
| IPSC input/output<br>curve<br>20 V | EAAT3glo<br>EAAT3glo/CaMKII | 614.60 ± 69.05<br>559.02 ± 101.72 | 7c/4a<br>7c/4a | P = 0.668 | Sup 2D |
| IPSC input/output<br>curve<br>25 V | EAAT3glo<br>EAAT3glo/CaMKII | 656.30 ± 84.60<br>590.20 ± 94.71 | 7c/4a<br>7c/4a | P = 0.616 | Sup 2D |
| IPSC input/output<br>curve<br>30 V | EAAT3glo<br>EAAT3glo/CaMKII | 717.92 ± 96.97<br>648.06 ± 106.64 | 7c/4a<br>7c/4a | P = 0.639 | Sup 2D |
| sIPSC<br>amplitude | EAAT3glo<br>EAAT3glo/GAD | 23.71 ± 2.11<br>28.16 ± 1.61 | 13c/8a<br>13c/8a | P = 0.107 | Sup. 3A |
| sIPSC<br>frequency | EAAT3glo<br>EAAT3glo/GAD | 11.09 ± 1.08<br>13.26 ± 1.55 | 13c/8a<br>13c/8a | P = 0.263 | Sup. 3A |
| mIPSC<br>amplitude | EAAT3glo<br>EAAT3glo/GAD | 15.01 ± 0.86<br>19.63 ± 0.98 | 7c/5a<br>7c/5a | P = 0.005 | Sup. 3B |
| mIPSC<br>frequency | EAAT3glo<br>EAAT3glo/GAD | 4.76 ± 0.86<br>7.26 ± 0.54 | 7c/5a<br>7c/5a | P = 0.015 | Sup. 3B |
| IPSC input/output<br>curve<br>5V | EAAT3glo<br>EAAT3glo/GAD | 400.33 ± 166.99<br>380.21 ± 36.26 | 5c/3a<br>5c/3a | P = 0.931 | Sup.3C |
| IPSC input/output<br>curve<br>10V | EAAT3glo<br>EAAT3glo/GAD | 555.89 ± 173.53<br>631.40 ± 122.81 | 5c/3a<br>5c/3a | P = 0.771 | Sup.3C |
| IPSC input/output<br>curve<br>15V | EAAT3glo<br>EAAT3glo/GAD | 637.56 ± 178.65<br>768.60 ± 93.78 | 5c/3a<br>5c/3a | P = 0.614 | Sup.3C |

| | | | | | |
|---|---|---|---|---|---|
| IPSC input/output curve 20V | EAAT3$^{glo}$<br>EAAT3$^{glo}$/GAD | 720.78 ± 188.17<br>886.60 ± 59.43 | 5c/3a<br>5c/3a | P = 0.537 | Sup.3C |
| IPSC input/output curve 25V | EAAT3$^{glo}$<br>EAAT3$^{glo}$/GAD | 746.44 ± 193.24<br>975.00 ± 37.27 | 5c/3a<br>5c/3a | P = 0.406 | Sup.3C |
| IPSC input/output curve 30V | EAAT3$^{glo}$<br>EAAT3$^{glo}$/GAD | 793.78 ± 206.40<br>1081.20 ± 76.49 | 5c/3a<br>5c/3a | P = 0.337 | Sup.3C |
| IPSC PPR 30 ms | EAAT3$^{glo}$<br>EAAT3$^{glo}$/GAD | 0.56 ± 0.06<br>0.55 ± 0.04 | 7c/4a<br>17c/9a | P = 0.841 | Sup.3D |
| IPSC PPR 70 ms | EAAT3$^{glo}$<br>EAAT3$^{glo}$/GAD | 0.53 ± 0.06<br>0.49 ± 0.03 | 7c/4a<br>17c/9a | P = 0.306 | Sup.3D |
| IPSC PPR 100 ms | EAAT3$^{glo}$<br>EAAT3$^{glo}$/GAD | 0.48 ± 0.05<br>0.48 ± 0.03 | 7c/4a<br>17c/9a | P = 0.469 | Sup.3D |
| IPSC PPR 300 ms | EAAT3$^{glo}$<br>EAAT3$^{glo}$/GAD | 0.51 ± 0.04<br>0.49 ± 0.03 | 7c/4a<br>17c/9a | P = 0.369 | Sup.3D |
| DHPG-induced depression of IPSCs | EAAT3$^{glo}$<br>EAAT3$^{glo}$/GAD | 37.53 ± 8.67<br>36.39 ± 7.13 | 5c/3a<br>4c/3a | P = 0.925 | Sup.3E |
| Glutamate transporter blockers on iLTD magnitude | EAAT3$^{glo}$ | 57.63 ± 5.07<br>TBOA 41.27 ± 4.85<br>INH3a 38.21 ± 7.19<br>DHKA 57.23 ± 8.39 | 9c/6a<br>7c/5a<br>7c/6a<br>7c/6a | <br>P = 0.040<br>P = 0.040<br>P = 0.965 | Sup.4A |
| fEPSP PP-LFS LTD | EAAT3$^{glo}$<br>EAAT3$^{glo}$/CaMKII | 57.58 ± 4.57<br>105.49 ± 11.64<br>INH3a 74.48 ± 5.63 | 8s/6a<br>10s/7a<br>8s/4a | P = 0.003<br>P = 0.003<br>P = 0.042 | Sup.4B |
